# Two telomere-to-telomere genome assemblies and comparisons revealed the conserved key genes associated with sugar accumulation in *Rubus* genus

**DOI:** 10.1101/2025.04.09.646780

**Authors:** Xiaopeng Li, Xianyan Han, Shoucheng Liu, Qunying Zhang, Jiyuan Guan, Yajun Tang, Meng Zhang, Pengbo Xu, Mengzhuo Zheng, Jianxin Bian, Kui Li, Guilian Sun, Yan Sun, Yuanxin Dong, Xiaojie Lin, Zhengdong Wang, Houcheng Zhou, Guohui Yang, Zhongchi Liu, Hongli Lian, Hang He, Junhui Zhou

## Abstract

For the first time, we assembled two highly continuous, completely gap-free reference genomes of the *Rubus* subgenus: *Rubus hirsutus* Thunb. ‘Penglei’ (XMM) and *Rubus eustephanos* Focke ex Diels ‘Dahongpao’ (DHP), which are widely distributed in southern China with similar phenotypic traits (Figures 1A, 1B, and Figure S1), yet ripe fruits display distinct sugar accumulation levels (Table S1), making them ideal candidates for investigating the mechanisms underlying sugar accumulation in the *Rubus* genus. The XMM (213.53 Mb, 28,204 genes) and DHP (218.26 Mb, 28,569 genes) genomes exhibit close evolutionary relationships, diverging approximately 3.21 Mya. Comparative genomics identified extensive synteny, interspecific structural variations (translocations, inversions, segmental duplications), and presence/absence variation (PAV). Using Hi-C interaction heatmaps and Sanger sequencing, we validated interspecific structural inversions. Additionally, we identified a sugar transporter gene (*MFS1*), which is present in XMM but absent in DHP. Combined analysis of the gene family expansion/contraction and transcriptome identified two conserved key genes (*RhSTP13* and *RhSTP7*) associated with sugar accumulation in *Rubus* genus and displayed distinct roles through transient expression assay. To facilitate functional genomics study, we also established a comprehensive *Rubus* database, RubusDB, a freely accessible repository consolidating all genomic, transcriptomic and phenotypic data of *Rubus* genus. These findings provide a foundational framework for elucidating the genetic basis of sugar accumulation, genome diversification, and trait improvement in *Rubus* species.

---

To the Editor,

The genus *Rubus*, commonly known as brambles, comprise small perennial berry shrubs within the Rosaceae family. These plants are composed of over 740 species and boast a rich diversity of germplasm resources with a cosmopolitan distribution spanning diverse latitudes [1, 2], rendering them of exceptional value for evolutionary studies. *Rubus* species, recognized by the United Nations Food and Agriculture Organization (FAO) as an international ‘third-generation fruit’, are particularly notable for their low sugar content and high abundance of bioactive compounds. These properties confer significant nutritional and medical value, including the ability to scavenge reactive oxygen species (ROS), reduce blood pressure, prevent carcinogenesis and cardiovascular diseases, and decelerate aging-related processes [3, 4]. Additionally, *Rubus* species exhibit a short juvenile phase, self-compatibility, and diploid genome (2n=2x=14), along with an facile transformation system [5], making an ideal model system for both fundamental and applied research in woody fruit crops.

Previously, the first assembled and annotated genome of *Rubus* species was black raspberry (*Rubus occidentalis* L.) [6], followed by the genomes of *Rubus chingii* Hu [7] and *Rubus idaeus* [8, 9]. The *Rubus* genus displays striking genomic and phenotypic diversity across its subgenera, a characteristic shaped by its perennial growth habit, high heterozygosity, and complex reproductive biology, resulting in substantial interspecific divergence. Besides, current *Rubus* genome assemblies remain fragmented, with substantial gaps in centromeric and telomeric regions and incomplete gene annotation. These limitations hinder comprehensive genomic analyses and functional studies. High-quality, telomere-to-telomere assemblies for representative subgenera are critically needed to advance *Rubus* research.

## RESULTS AND DISCUSSION

### Sequencing, assembly, and annotation of the XMM and DHP genome

The telomere-to-telomere (T2T) genomes of XMM and DHP were assembled using an integrative approach that combined ultra-long reads from Oxford Nanopore Technology (ONT), PacBio high-fidelity (HiFi) long reads, Illumina paired-end short reads, and High-throughput Chromosome Conformation Capture (Hi-C) data (Figure 1C, Figures S2, S3, and Table S2). This effort resulted in two of the most complete *Rubus* genomes to date (Table S3), featuring fully resolved telomeres enriched with the plant telomeric repeat motif (TTTAGGG) (Table S4). The genome sizes are 213.53 Mb for XMM and 218.26 Mb for DHP (Figure 1D and Table S5). Both assemblies achieved exceptional BUSCO completeness scores of 99.18% for XMM and 99.23% for DHP. Furthermore, the mapping rates for HiFi and ONT reads were 99.90% and 99.99% for XMM (with 100% coverage each) and 99.17% and 99.98% for DHP (with coverage of 99.87% and 99.94%, respectively). The LTR Assembly Index (LAI) values exceeded gold standards, with scores of 26.49 for XMM and 20.53 for DHP, while consensus quality value (QV) scores ranged from 65 to 68 (Table S6). Additionally, we identified two types of centromeric satellites (CEN18 and CEN17), which share 84% sequence similarity and occupy 19.94% and 23.90% of the centromeric regions in XMM and DHP, respectively (Figure 1D, Figure S4, and Table S7).

**FIGURE 1.**
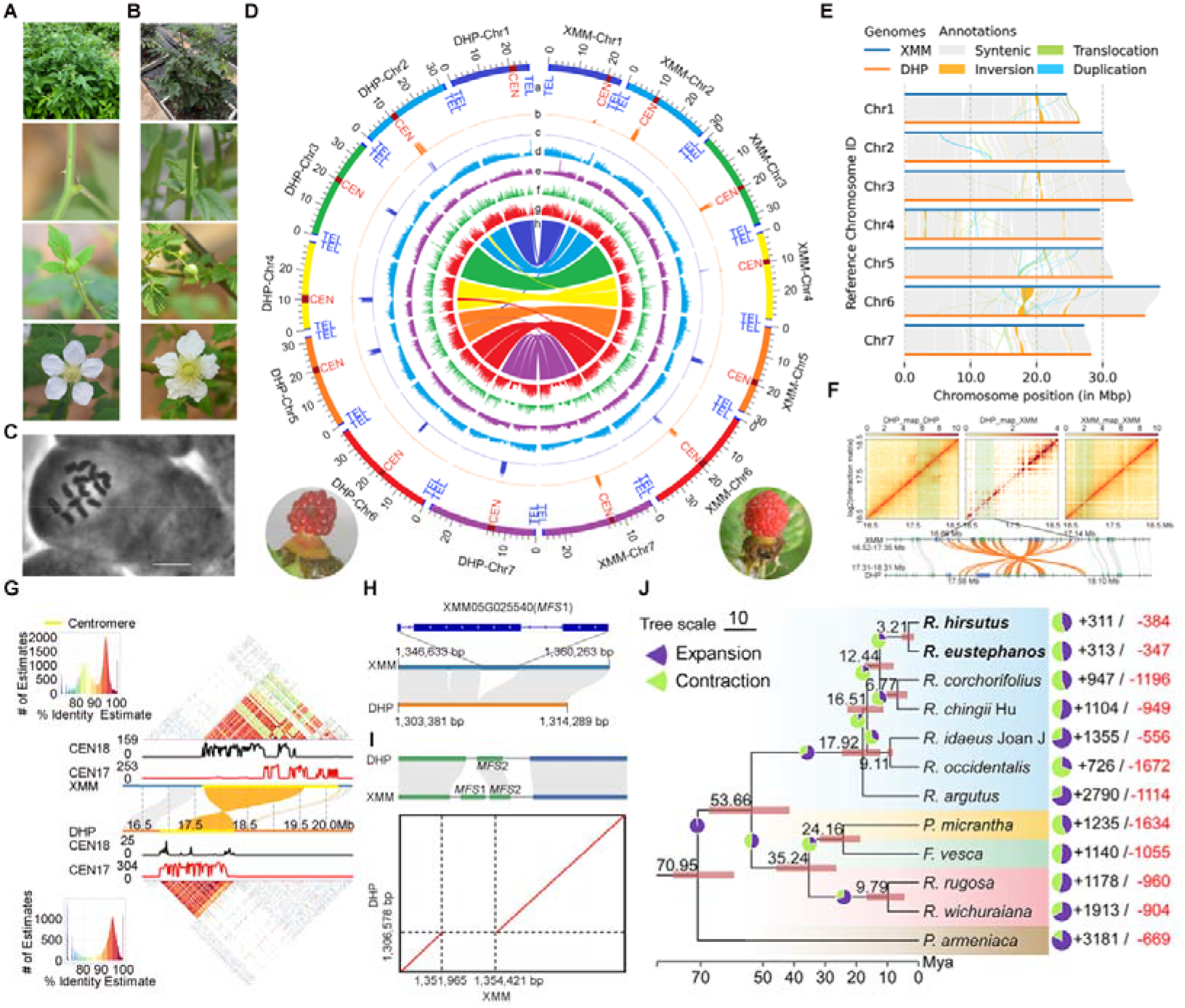
Phenotypic characterization, *de novo* genome assemblies, variation, and evolution of *Rubus hirsutus* Thunb. ‘Penglei’ (XMM) and *Rubus eustephanos* Focke ex Diels ‘Dahongpao’ (DHP). (A) Illustration of XMM whole plants, stem, flower bud and flower, respectively. (B) Illustration of DHP whole plants, stem, flower bud and flower, respectively. (C) The chromosome karyotype analysis of XMM. Bar = 10 µm. (D) Circular diagram of XMM and DHP reference genomes. (a) Chromosomes are represented by centromeres (dark red) and telomeres (blue); (b) CEN18 density; (c) CEN17 density; (d) Gene density; (e) TE density; (f) PAV density; (g) SNP density; (h) Collinear lines at the center of diagram highlight homoeologous chromosomes relationships and non-homoeologous regions. (E) Genomic alignments between XMM and DHP. Inversions, duplications and translocations are marked with orange, blue and green ribbons, respectively. (F) Identification of large inversions in chromosome 7 between XMM and DHP. The three heatmaps show the chromatin interaction matrix, including mapping Hi-C data of DHP against DHP genome (left), mapping Hi-C data of DHP against XMM genome (middle) and mapping Hi-C data of XMM against XMM genome (right). The lower panel illustrates gene alignments between XMM and DHP. (G) Synteny, structural rearrangements, monomer, and identity distribution in the 16-20 Mbp region of chromosome 6 between XMM and DHP. (H) Local genome synteny of chromosome 5, with a structural variation that is present only in XMM. The PAV region includes a gene related to sugar transporter (*MFS1*, XMM05G025540). (I) Gene-level matchings of *MFS* genes in the PAVs region between XMM and DHP. (J) Phylogenetic relationships and divergence times between raspberries and other Rosaceae species. The black numbers close to the divergence nodes indicate the divergence times and the red bars represent the 95% confidence intervals. The red and black numbers indicate expanded and contracted ortholog groups at the corresponding node. Scale bar corresponds to 10 Mya.

Through a combination of *ab initio* and homology-based approaches, 46.59% (99.48 Mb) and 47.18% (102.98 Mb) of repetitive sequences were identified in the XMM and DHP genomes, respectively (Table S8). Following repeat masking, gene structure annotation was performed using a hybrid strategy that integrated *de novo* prediction, homology-based methods, and transcriptome evidence (Figure S5). This comprehensive annotation revealed 28,204 and 28,569 protein-coding genes in the XMM and DHP genomes, respectively (Table S9). These annotations achieved exceptional coverage of 98.50% and 98.45% of the complete BUSCO gene set, surpassing the quality of all previously released *Rubus* genomes (Table S10).

### Comparative genomic analyses between XMM and DHP

To explore chromosomal variations and trait diversity between the XMM and DHP genomes, we conducted a comparative genomic analysis (Figures 1D, 1E). This analysis identified 380 syntenic blocks with over 79.00% 1-to-1 collinearity, encompassing 939,717 single nucleotide polymorphisms (SNPs), 180,328 insertions/deletions (InDels), and 867 structural variants, including 349 translocations, 408 duplications, and 110 inversions (Table S11). Notably, a large inversion on chromosome 7, containing 33 annotated genes, was confirmed by aligning Hi-C reads from DHP to the XMM genome and visualizing the Hi-C interaction heatmap (Figure 1F). Furthermore, a significant inversion was observed specifically in the centromeric region of chromosome 6 in both XMM and DHP genomes, which was validated through Sanger sequencing (Figures 1E, 1G, Figure S6, and Table S12).

Additionally, we identified 2,013 and 2,146 presence/absence variation (PAV) sequences in the XMM and DHP genomes, respectively, totaling 1.33 Mb and 1.28 Mb. Over 87.50% of these PAV regions were shorter than 1 kb (Figure S7A). Of particular interest, a sugar transporter gene (*MFS1*), present in the XMM genome but absent in DHP, exhibited higher expression levels across six tissues in XMM (Figures 1D, 1H, 1I and Figure S7B).

### Gene family expansion/contraction analysis and divergence time estimation

To explore the evolutionary history of the two *Rubus* species, we conducted a phylogenetic analysis using ten other published Rosaceae genomes (*Prunus armeniaca, Potentilla micrantha, Fragaria vesca, Rosa rugosa, Rosa wichuraiana, Rubus occidentalis, Rubus idaeus* ‘Joan J’, *Rubus argutus, Rubus chingii* Hu, and *Rubus corchorifolius*) from the Genome Database for Rosaceae [10]. Based on sequence similarity, we identified 33,275 ortholog groups, encompassing 27,209 genes from XMM and 27,527 genes from DHP (Figure 1J). A phylogenetic tree was constructed for these 12 Rosaceae species using 3,969 conserved single-copy genes. The analysis revealed that XMM and DHP form a distinct clade, having diverged from the common ancestor of four *Rubus* species approximately 12.44 million years ago (Mya). The divergence between XMM and DHP occurred more recently, around 3.21 Mya (Figure S8). Additionally, *Rubus chingii* Hu and *Rubus corchorifolius* were found to be closely related to XMM and DHP.

Following the divergence of XMM and DHP, 198 gene families underwent significant expansion, while 617 families experienced contraction. Notably, gene families associated with glycoprotein biosynthetic and metabolic process expanded, whereas those involved in (1->3)-beta-D-glucan biosynthetic contracted (corrected *P* < 0.05) (Figures S9, S10). Further analysis of expanded and contracted gene families, along with functional gene annotations, revealed that two sugar-related gene families in XMM have undergone expansion, while three sugar-related gene families have undergone contraction (Figure 2A and Table S13).

**FIGURE 2.**
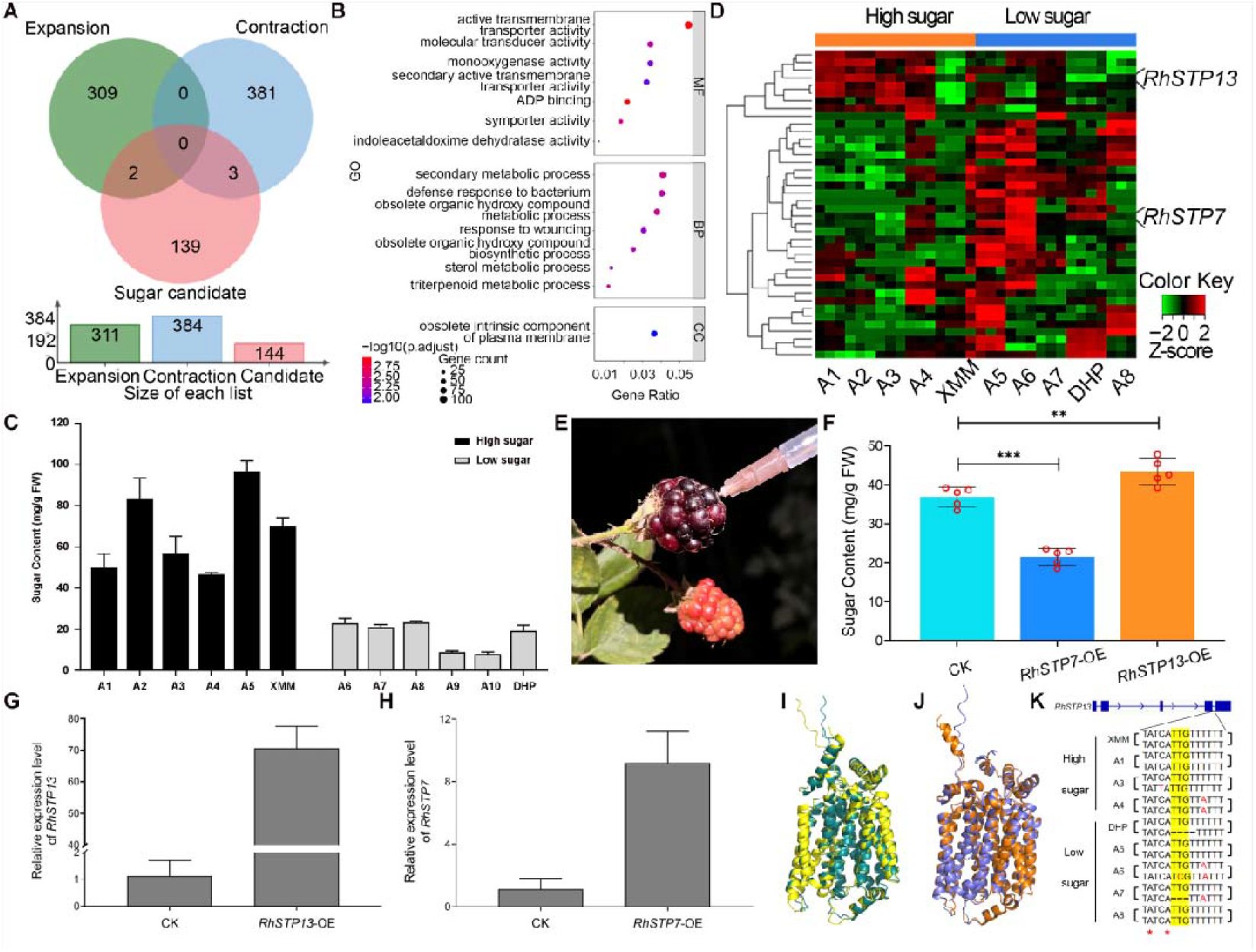
Identification and functional verification of candidate genes for sugar accumulation in *Rubus*. (A) Venn diagram depicts the family of sugar-related, contraction and expansion gene family in XMM. (B) GO enrichment analyses of DEGs in ripe fruits between high-sugar and low-sugar groups. (C) Total sugar contents of ripe fruits in ten *Rubus* species were detected through UPLC-MS/MS platform. A1-A8 represented *Rubus idaeus* Tulamen, *Rubus idaeus* Erika, *Rubus idaeus* Orange Miracle, *Rubus chingii* Hu, *Rubus setchuenensis* Bureau et Franch, *Rubus argutus* Hillquist, *Rubus idaeus* Heritage and *Rubus occidentalis*, respectively. (D)Sugar-relate allelic expression profiles between five high-sugar and five low-sugar samples. A1-A8 represented *Rubus idaeus* Tulamen, *Rubus idaeus* Erika, *Rubus idaeus* Orange Miracle, *Rubus chingii* Hu, *Rubus setchuenensis* Bureau et Franch, *Rubus argutus* Hillquist, *Rubus idaeus* Heritage and *Rubus occidentalis*, respectively. The color gradient ranging from red to green is indicative of the relative expression values of the genes. (E) Over-expression constructions were carried out transient transformation in *Rubus argutus* fruits at turning stage. (F) Total sugar content in mature fruits of *RhSTP7*-OE and *RhSTP13*-OE based on UPLC-MS/MS results. Cas9-OE infiltrated-fruits were used as control. Two biological replicates consisting of five fruits were prepared for determination of sugar content. (G-H) Relative expression level of two candidate genes in agro-infiltrated *Rubus argutus* fruits, *RhSTP13*-OE (G) and *RhSTP7*-OE (H) construction. (I-J) Protein structural prediction of *RhSTP13* (I) and *RhSTP7* (J). Deep teal and orange represent *RhSTP13* and *RhSTP7* in XMM. Yellow and slate represent allele in DHP. (K) Multiple sequence alignment of the fourth intron region of *RhSTP13* in different *Rubus* genus for Illumina sequencing data. ‘*’ represented potential branch point.

### Identifying candidate genes of sugar accumulation in *Rubus* fruits

Based on our UPLC-MS/MS results (Figure 2C), XMM exhibited a remarkably high sugar content, approximately 3.64 times greater than that of DHP. To broaden our analysis, we carried out transcriptome sequencing and annotated eight other *Rubus* materials. These 10 materials were categorized into high-sugar and low-sugar groups according to their total fruit sugar content (Table S14), revealing 3,531 differentially expressed genes (DEGs) which were significantly enriched in the molecular function ‘active transmembrane transporter activity’ (GO:0022804) (Figure 2B and Table S15). We further identified 40 candidate genes primarily involved in sugar transport processes that likely contribute to the differences in sugar accumulation between XMM and DHP by integrating differential gene expression analysis, homologous gene comparison, and functional annotation (Figure 2D and Table S16).

### *RhSTP13 and RhSTP7* exhibit distinct regulatory effects on sugar content in

#### Rubus

Then we selected two most striking DEGs, *RhSTP13* and *RhSTP7*, which were enriched in the ‘active transmembrane transporter activity’ and ‘symporter activity’ GO terms and exhibited highest contrasting expression patterns: *RhSTP13* was upregulated in high-sugar samples, while *RhSTP7* was downregulated. To verify their functions in sugar accumulation during *Rubus* fruit development, we constructed *RhSTP13-* and *RhSTP7-*overexpressed constructs and performed transient transformation assays in the fruits of *Rubus argutus*, a cultivar with medium sugar content (Figure 2E). Overexpression of *RhSTP13* led to a 17.48% increase in sugar concentration, whereas overexpression of *RhSTP7* resulted in a 41.73% decrease in total sugar content (Figure 2F, and Table S17). The relative expression levels of *RhSTP13* and *RhSTP7* were consistent to the expected changes in sugar content (Figures 2G, 2H). In addition, we predicted the protein 3D structures of both genes and found no significant structural differences (Figures 2I, 2J). *RhSTP13* belongs to an expanded gene family in XMM. We investigated the conserved SV sequences via *Rubus* resequencing and Sanger sequencing, and identified multiple mutation types within TTG sequence near to the branch point in the fourth intron region of *RhSTP13* in most low-sugar groups. This mutation may contribute to variations in *RhSTP13* expression levels, ultimately influencing sugar accumulation (Figures 2D, 2K and Figure S11). These T2T genomes provide a robust foundation for genomic and functional studies of *Rubus* genus, particularly in understanding the genetic basis of sugar accumulation and other economical traits.

### Establishment of RubusDB database

We have established a comprehensive database for the *Rubus* genus, RubusDB, to document all genomic, transcriptomic, and phenotypic data generated in this study (Figure S12). Additionally, RubusDB offers basic analysis tools, such as batch query and BLAST, to support gene discovery and genomic investigations across all species within the *Rubus* genus. We will continue incorporate newly-updated genomic resources and bioinformatics tools of all *Rubus* genus to RubusDB in the future.

## METHODS

Comprehensive detailed procedures are provided in the Supplementary Information.

## AUTHOR CONTRIBUTIONS

**Xiaopeng Li:** Investigation; methodology; formal analysis; writing-review and editing. **Xianyan Han:** Investigation; methodology; writing-review and editing. **Shoucheng Liu:** Investigation; formal analysis; writing-review and editing. **Qunying Zhang:** Funding acquisition; resources; investigation. **Jiyuan Guan:** Methodology; resources. **Yajun Tang:** Methodology; validation. **Meng Zhang:** Investigation. **Pengbo Xu:** Investigation. **Mengzhuo Zheng:** Formal analysis; writing-review and editing. **Jianxin Bian:** Investigation; formal analysis. **Kui Li:** Investigation; writing-review and editing. **Guilian Sun:** Methodology; investigation. **Yan Sun:** Investigation. **Yuanxin Dong:** Investigation. **Xiaojie Lin:** Investigation. **Zhengdong Wang:** Validation. **Houcheng Zhou:** Methodology; supervision. **Guohui Yang:** Resources. **Zhongchi Liu:** Project administration; conceptualization. **Hongli Lian:** Project administration; supervision; writing-review and editing. **Hang He:** Project administration; conceptualization; supervision; writing-review and editing. **Junhui Zhou:** Project administration; funding acquisition; conceptualization; resources; supervision; writing-review and editing.

## ACKNOWLEDGEMENT

We thank the high-performance computing platform, Dr. Bosheng Li, Dr. Guochen Qin, Dr. Xiaoli Lin, Zhiying Lou, Dr. Dongdong Lu, Bailong Song and Hongxiao Sun (PKU-IAAS) for technical assistance in this project. We also thank Taishan Scholars Program (J.Z.), the Dean’s Foundation from PKU-IAAS, the funding from Shandong Laboratory of Advanced Agricultural Sciences at Weifang, and Guizhou Provincial Scientific and Technological Program (QKHFQ[2024]004-1) for financial support.

## DATA AVAILABILITY STATEMENT

All the raw sequencing data generated for this project, including the Nanopore ONT data, PacBio HiFi data, and Illumina NovaSeq data, as well as the assemblies, are archived at the National Genomics Data Center.

## ETHICS STATEMENT

No animals or humans were involved in this study.

## SUPPLEMENTAL INFORMATION

Additional supporting information can be found online in the Supporting Information section at the end of this article.

**Figure S1**. Morphological comparison between *Rubus hirsutus* Thunb. ‘Penglei’ (XMM) and *Rubus eustephanos* Focke ex Diels ‘Dahongpao’ (DHP).

**Figure S2**. Ploidy characterization of XMM and DHP.

**Figure S3**. Assembly pipeline for *Rubus* genome.

**Figure S4**. Representative pairwise sequence identity heatmap for centromeres located on XMM chromosomes and DHP chromosomes.

**Figure S5**. Pipeline of genome annotation for XMM and DHP.

**Figure S6**. Structural variants between XMM and DHP genomes.

**Figure S7**. PAV regions of XMM and DHP.

**Figure S8**. Phylogenetic relationships between *Rubus* genus and other Rosaceae species.

**Figure S9**. GO enrichment analyses of significant expansion gene families during the divergence of XMM and DHP.

**Figure S10**. GO enrichment analyses of significant contraction gene families during the divergence of XMM and DHP.

**Figure S11**. Multiple sequence alignment of the fourth intron region of *RhSTP13* in different *Rubus* genus.

**Figure S12**. Overview of data sources, tools, and module interactions of RubusDB.

**Table S1**. Total sugar content, total acidity and sugar-acid ratio of *Rubus hirsutus* Thunb. ‘Penglei’ (XMM) and *Rubus eustephanos* Focke ex Diels ‘Dahongpao’ (DHP)

**Table S2**. Data used for *Rubus hirsutus* Thunb. ‘Penglei’ (XMM) and *Rubus eustephanos* Focke ex Diels ‘Dahongpao’ (DHP) assembly and annotation.

**Table S3**. Statistics of the published Rubus genomes (all values are anchored to the chromosomes).

**Table S4**. Telomere statistics of XMM and DHP genome assemblies.

**Table S5**. Length statistics of XMM and DHP genome assemblies.

**Table S6**. Evaluation of XMM and DHP genome assemblies.

**Table S7**. Centromere statistics of XMM and DHP genome assemblies.

**Table S8**. Summary of XMM and DHP TE assemblies (bp).

**Table S9**. Summary of XMM and DHP gene annotations.

**Table S10**. BUSCO analysis of published *Rubus* genome annotation.

**Table S11**. Summary of aligned sequences and structural variations in the XMM and DHP genomes.

**Table S12**. Primers used in this study.

**Table S13**. Contraction and expansion of sugar-related gene families in XMM.

**Table S14**. Total sugar contents of ripe fruits of ten *Rubus* species were detected using UPLC-MS/MS platform.

**Table S15**. The enriched GO terms for DEGs in XMM.

**Table S16**. 40 sugar-related candidate genes.

**Table S17**. Total sugar content of *RhSTP7*-OE, *RhSTP13*-OE fruits and control.

