## Supplemental information for "Two telomere-to-telomere genome assemblies and comparisons revealed the conserved key genes associated with sugar accumulation in *Rubus* genus" (from another version of this preprint: version 1, 2025-04-10)

**Contents**

**Supplemental figures S1-S11**

**Supplemental materials and methods**

**Supplemental references**

### Supplemental figures

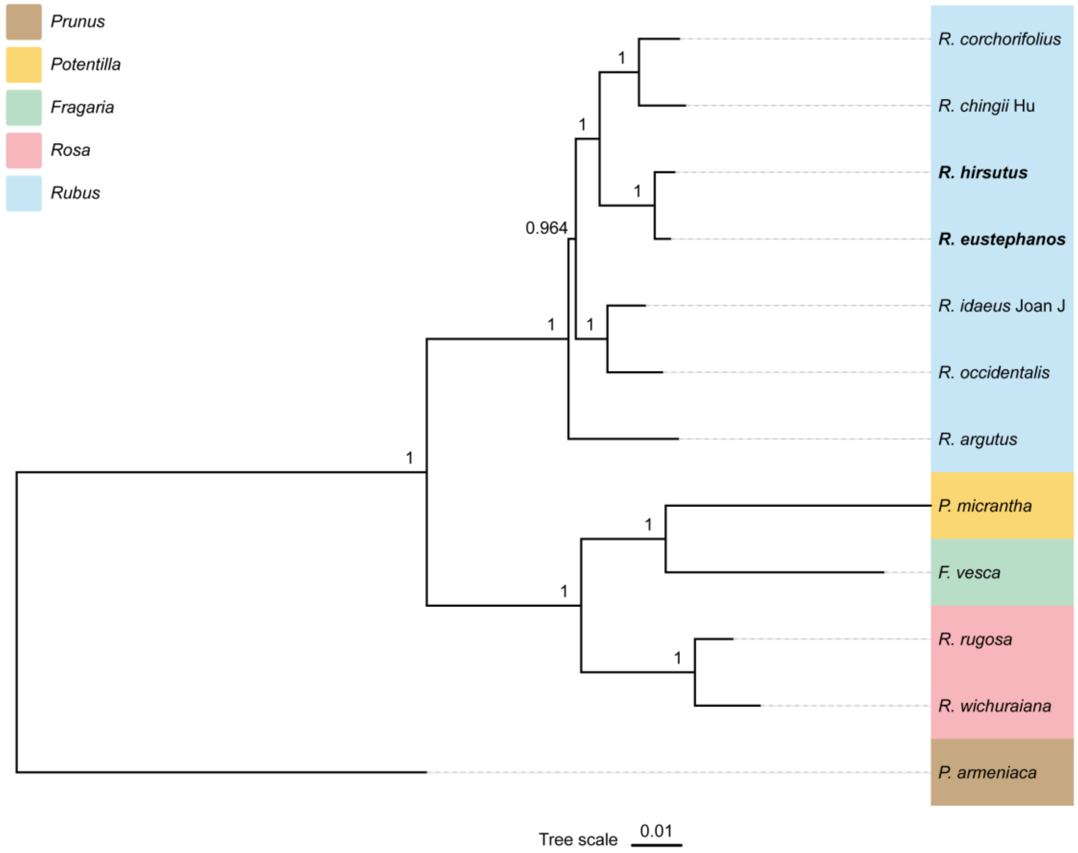


**Supplemental Figure 1. Phylogenetic relationships between *Rubus* genus and other Rosaceae species.**

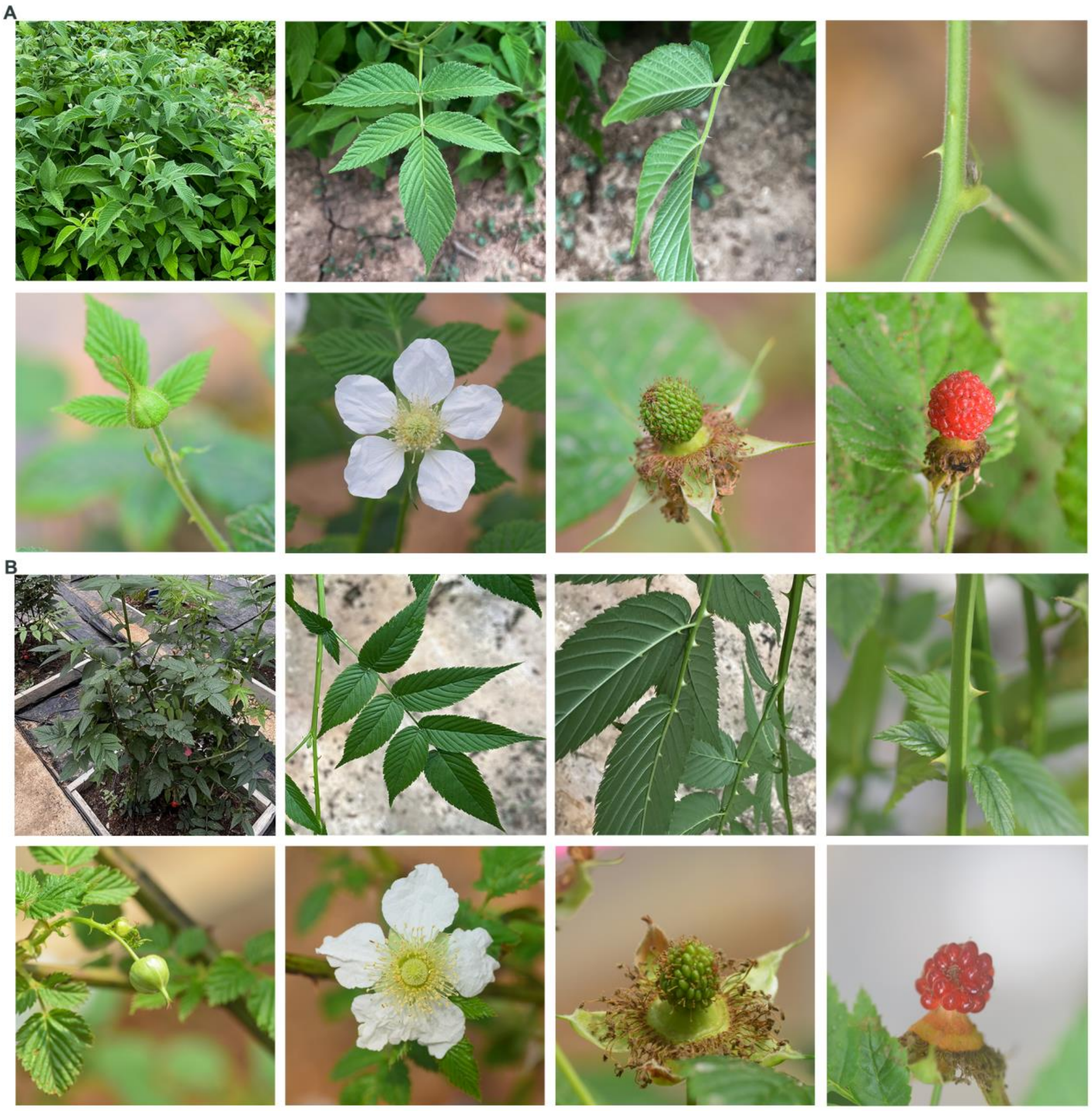


**Supplemental Figure 2. Morphological Comparison between *Rubus hirsutus* Thunb. 'Penglei' (XMM) (A) and *Rubus eustephanos* Focke ex Diels 'Dahongpao' (DHP) (B).** Illustration of whole plants, leaves (front), leaves (back), stem, flower bud, flower, unripe fruit, and ripe fruit, respectively.

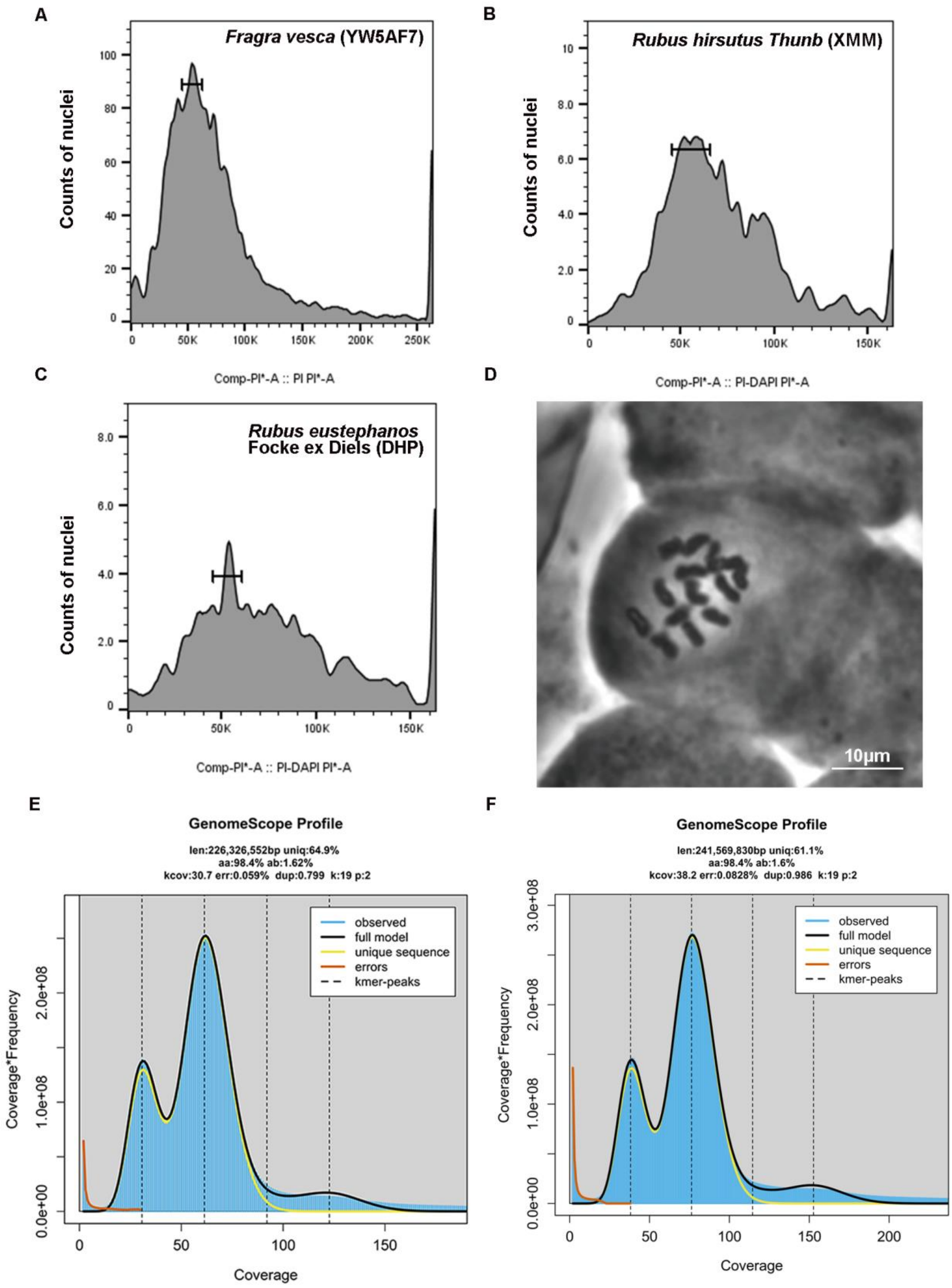


**Supplemental Figure 3. Ploidy detection of XMM and DHP. (A)** *Fragra vesca* YW5AF7 (2n=2x=14) was used as a reference standard of diploid by flow cytometry. **(B-C)** Flow cytometry analysis of the ploidy identifications of both XMM **(B)** and DHP **(C)** are diploid. **(D)** The chromosome karyotype analysis of XMM. **(E-F)** K-mer distribution and statistics for the genome of XMM **(E)** and DHP **(F)**.

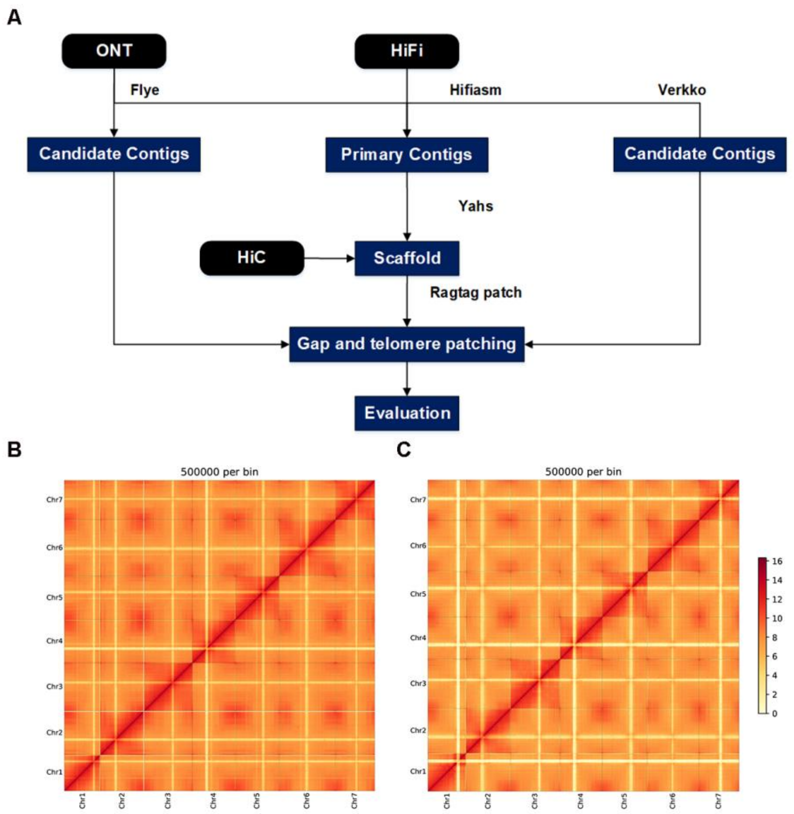


**Supplemental Figure 4. Two complete genome assemblies of XMM and DHP.**

**(A)** Pipeline of assembling XMM and DHP. **(B-C)** Intensity signal heat map of Hi-C chromosome interaction of XMM **(B)** and DHP **(C)**.

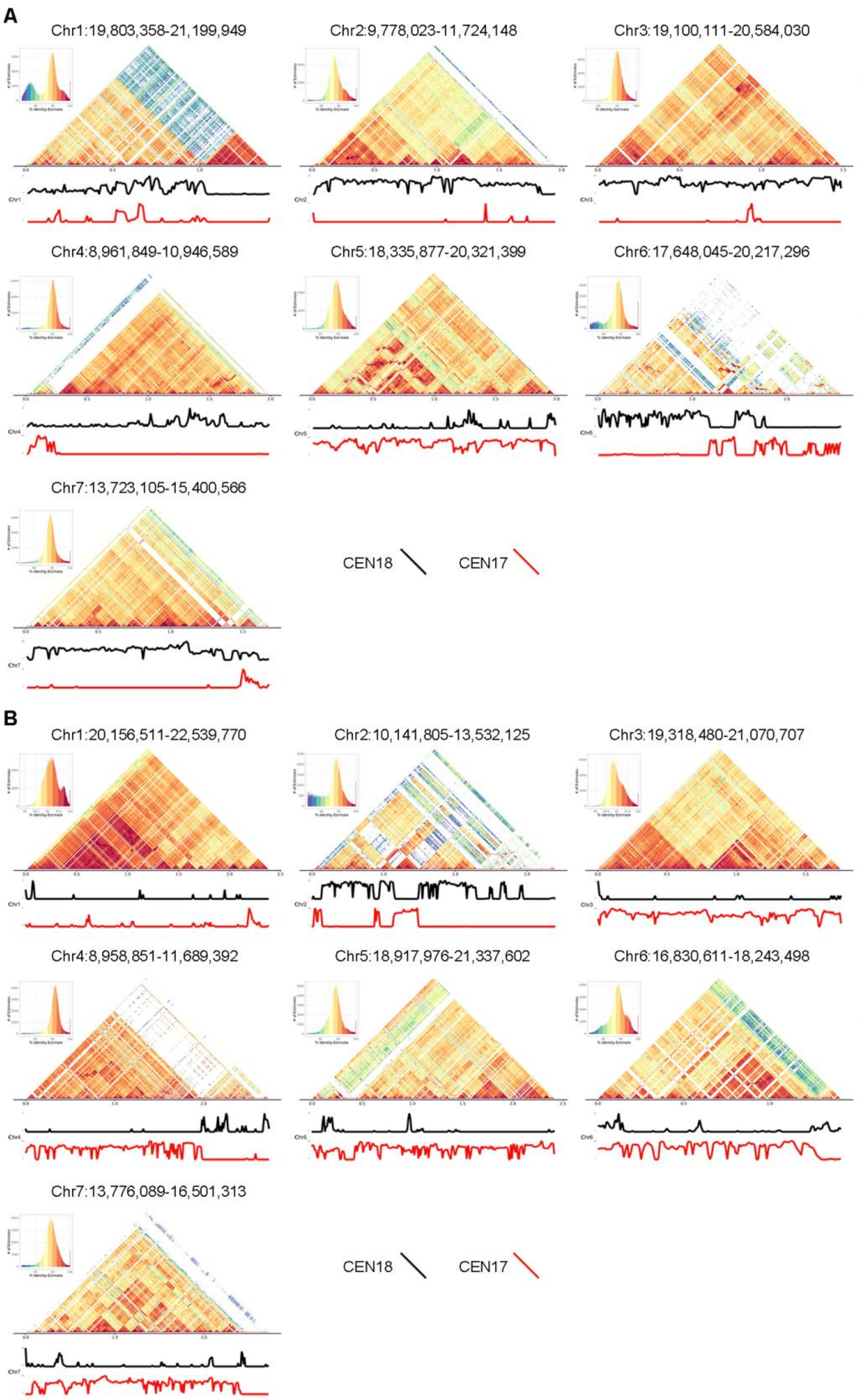


**Supplemental Figure 5. Representative pairwise sequence identity heatmap for centromeres located on XMM chromosomes (A) and DHP chromosomes (B).**

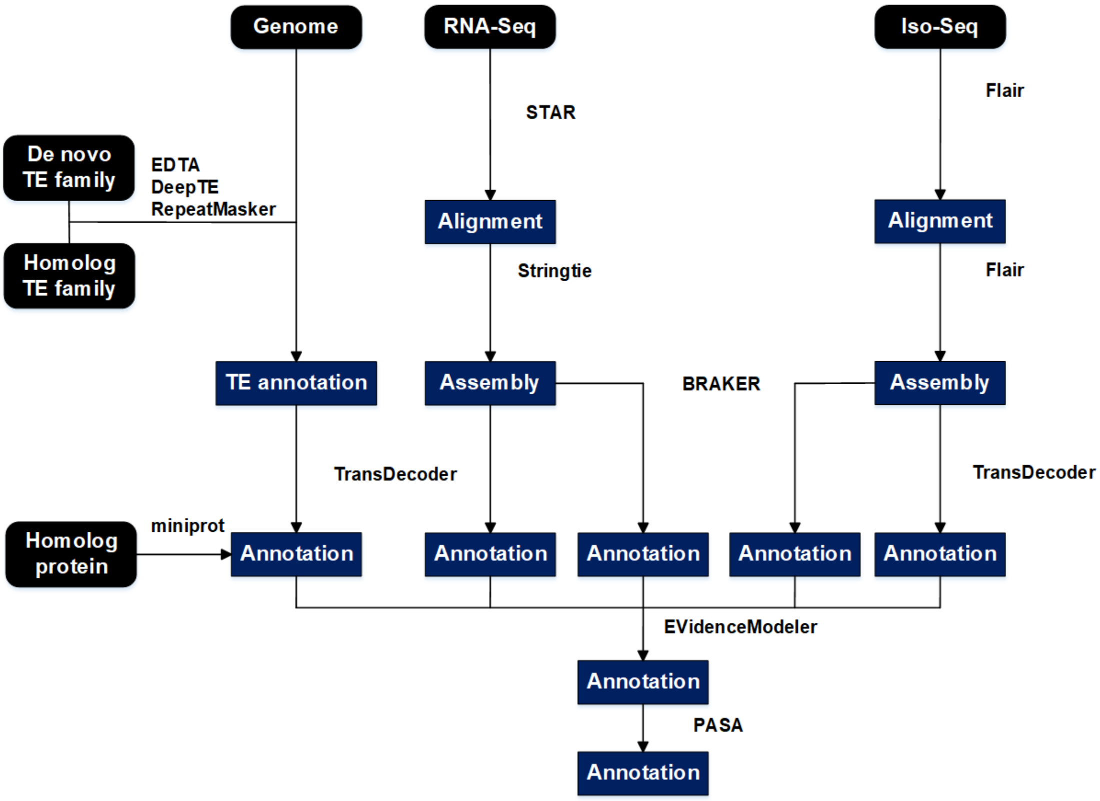


**Supplemental Figure 6. Pipeline of genome annotation for XMM and DHP.**

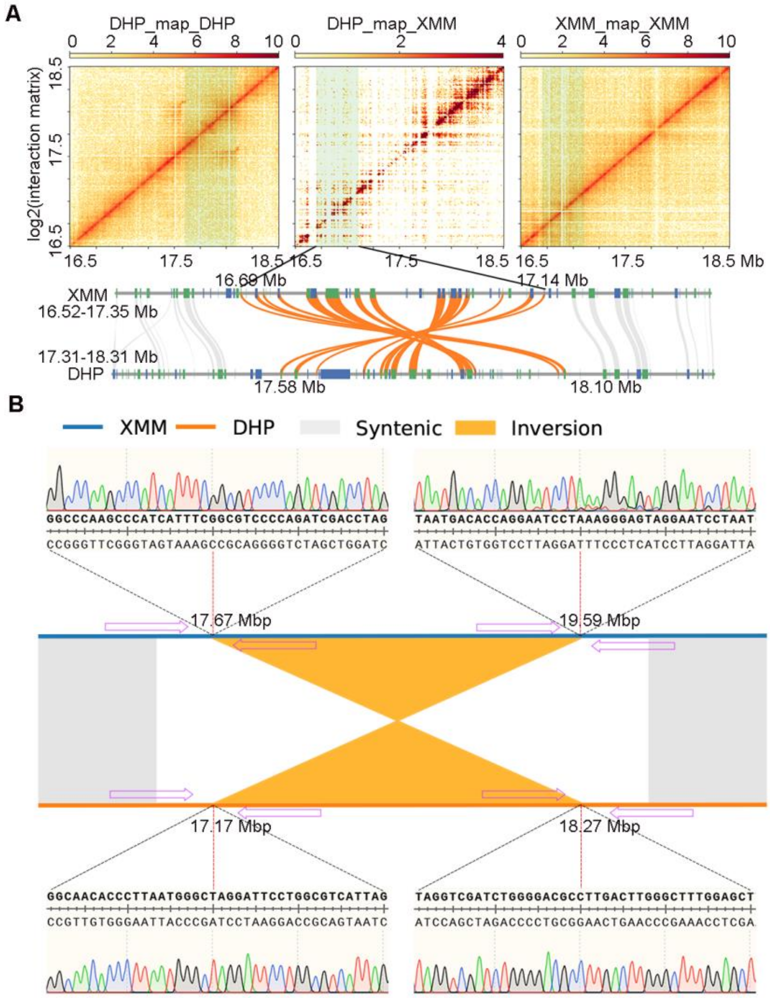


**Supplemental Figure 7. Structural variants between XMM and DHP genomes. (A)** Identification of large inversions in chromosome 7 between XMM and DHP. The three heatmaps show the chromatin interaction matrix, including mapping Hi-C data of DHP against DHP genome (left), mapping Hi-C data of DHP against XMM genome (middle) and mapping Hi-C data of XMM against XMM genome (right). The lower panel illustrates gene alignments between XMM and DHP. **(B)** Sanger sequencing combined with PCR validated the chromosome 6 centromeric inversion. The figure shows the sequences of four pairs of primers and partial Sanger sequencing (Supplemental table 12) results at the inversion boundaries. The sequence shown is the Sanger sequencing result for this location.

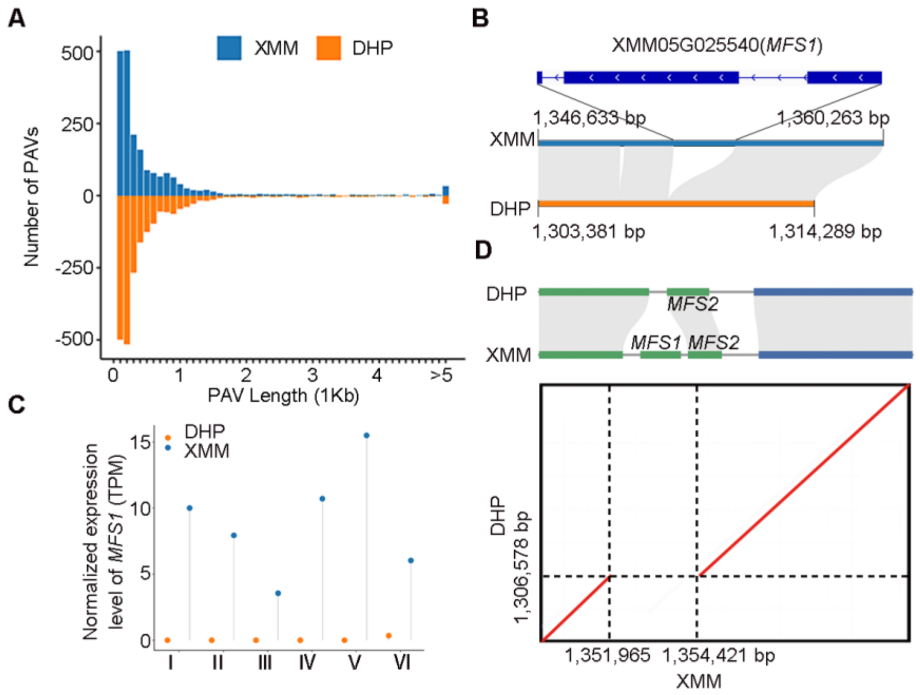


**Supplemental Figure 8. PAV regions of XMM and DHP. (A)** Distribution of lengths for PAVs in XMM and DHP. **(B)** Local genome synteny of chromosome 5, with a structural variation that is present only in XMM. The PAV region includes a gene related to sugar transporter (*MFS1*, XMM05G025540). **(C)** Normalized expression level of *MFS1* in six different tissues of XMM and DHP. Stages I-VI indicate the tissues of root, stem, leaf, flower, unripe fruits, and ripe fruits, respectively. **(D)** Gene-level matchings of *MFS* genes in the PAVs region between XMM and DHP.

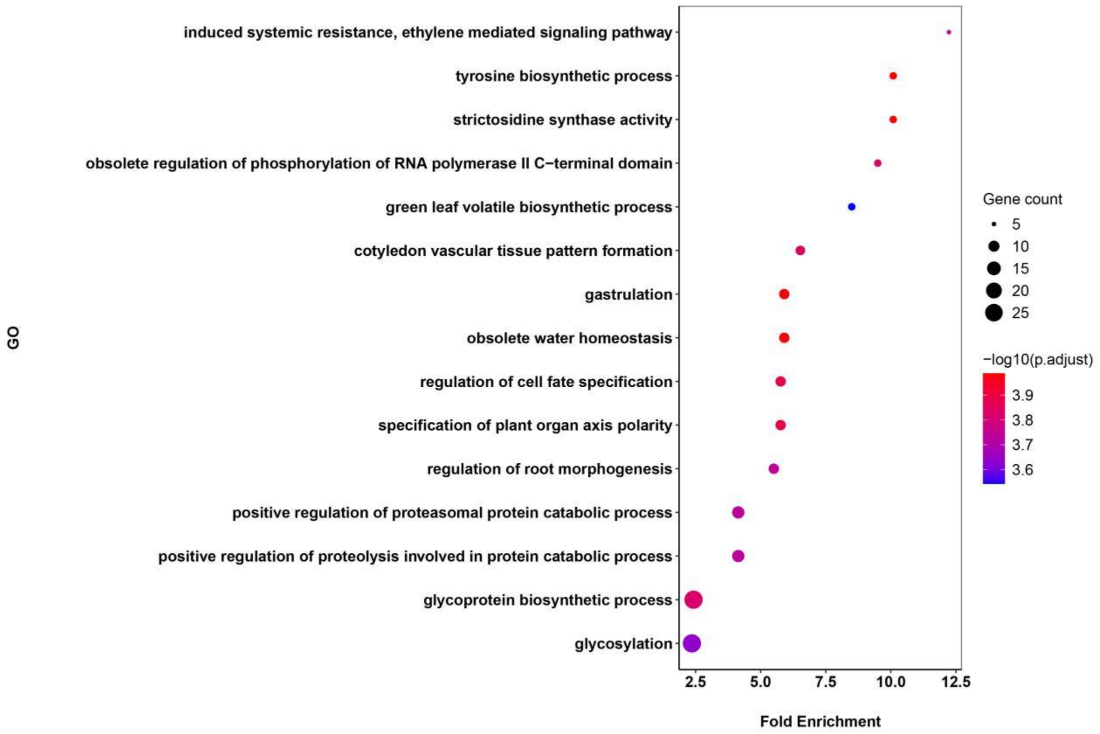


**Supplemental Figure 9. GO enrichment analyses of significant expansion gene families during the divergence of XMM and DHP.**

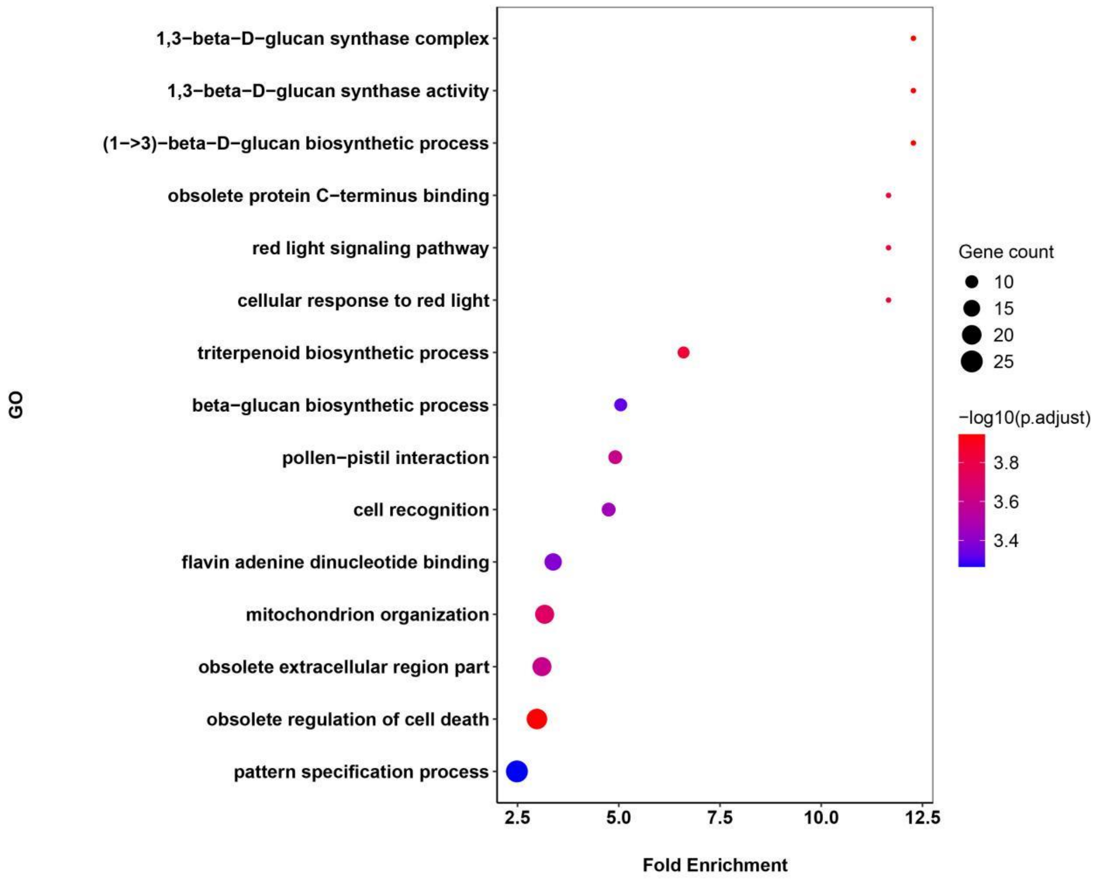


**Supplemental Figure 10. GO enrichment analyses of significant contraction gene families during the divergence of XMM and DHP.**

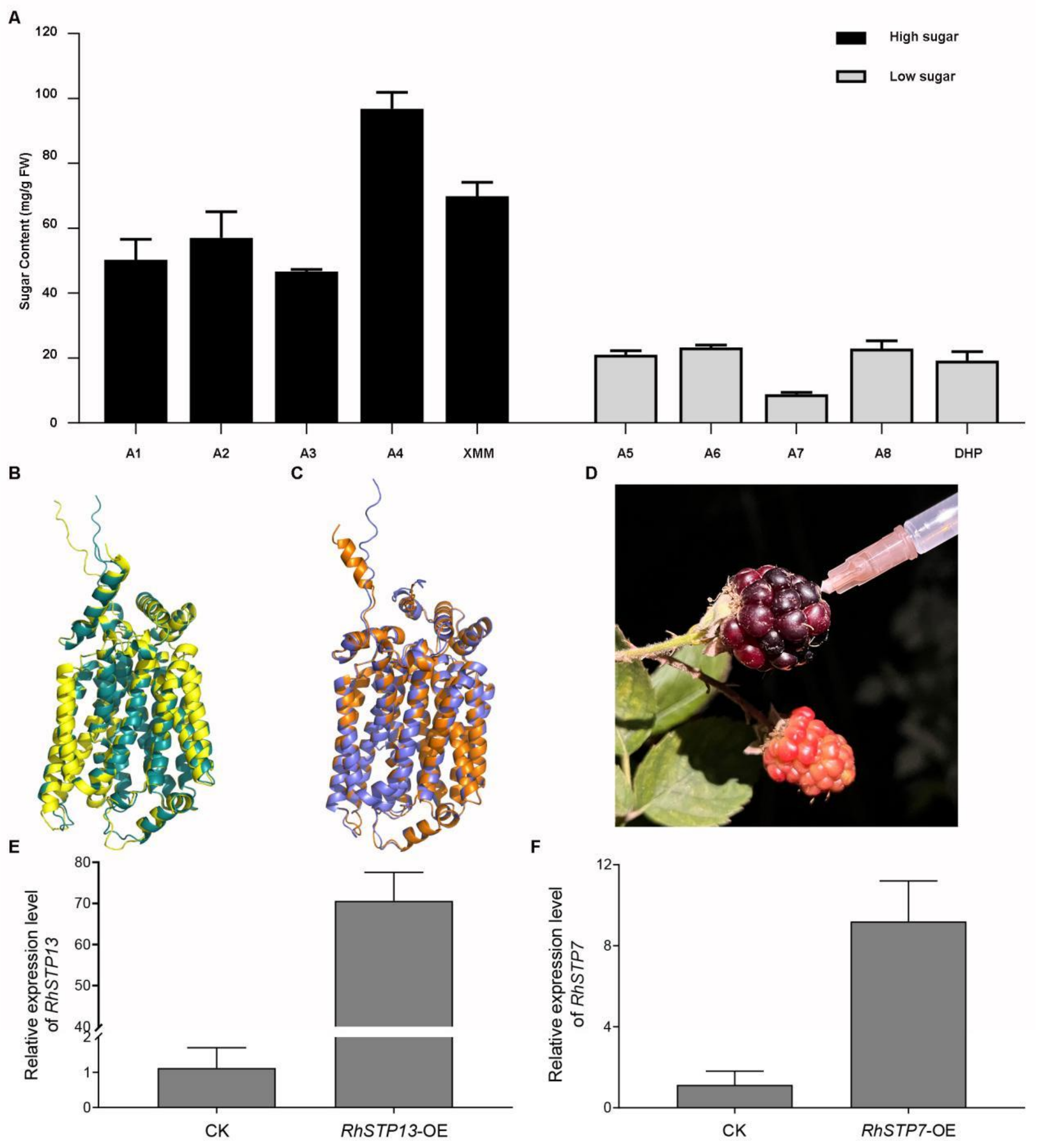


**Supplemental Figure 11. Functional verification of *RhSTP13* and *RhSTP7*.**

**(A)** Total sugar contents of ripe fruits in ten *Rubus* species were detected through UPLC-MS/MS platform. A1-A8 represented *Rubus idaeus* Tulamen, *Rubus idaeus* Erika, *Rubus idaeus* Orange Miracle, *Rubus chingii* Hu, *Rubus setchuenensis* Bureau et Franch, *Rubus argutus* Hillquist, *Rubus idaeus* Heritage and *Rubus occidentalis*, respectively. **(B-C)** Protein structural prediction of *RhSTP13* **(B)** and *RhSTP7* **(C)**. Deep teal and orange represent *RhSTP13* and *RhSTP7* in XMM. Yellow and slate represent allele in DHP. **(D)** Over-expression constructions were carried out transient transformation in *Rubus argutus* fruits at turning stage. **(E-F)** Relative expression level of two candidate genes in agro-infiltrated *Rubus argutus* fruits, *RhSTP13*-OE **(E)** and *RhSTP7*-OE **(F)** construction.

### Supplemental materials and methods

#### Plant materials

The individual *Rubus hirsutus* Thunb. 'Penglei' (XMM) and *Rubus eustephanos* Focke ex Diels 'Dahongpao' (DHP) plants planted in a raspberry germplasm nursery which located in Guizhou Botanical Garden (106°43'E, 26°37'N) were used for genome and transcriptome experiments.

#### Genomic DNA extraction, library construction, and sequencing

Tender leaves of XMM and DHP were washed with sterile water, frozen immediately with liquid nitrogen, and stored at -80℃. Genomic DNA (gDNA) was extracted from the young leaf of XMM and DHP using a DNAsecure Plant Kit (TIANGEN, DP320). High quality gDNA (≥10 μg, ≥100 ng/μL) was used to construct a HiFi SMRTbell library using the SMRTbell® Express Template Prep Kit 3.0. The Hi-C library was constructed as follows: crosslinking cells with formaldehyde, digesting DNA with a suitable 4-cutter restriction enzyme *Mbo*I, filling ends and mark with biotin, ligating the resulting blunt-end fragments, purification and random shearing DNA into 300-500 bp fragments. Paired-end sequencing with 150 base pairs (bp) reads was conducted on the Illumina Novaseq 6000 platform (Berry Genomics Co., Ltd). The paired-end library, featuring a fragment size of 350 bp, was meticulously prepared following the manufacturer's guidelines (Illumina) and subsequently sequenced on the Illumina Novaseq 6000 platform. The library was basically prepared by following steps: DNA fragmentation by sonication, end-polishing of the DNA fragments, A-tailing and ligation with the full-length adapters for Illumina sequencing, PCR amplification, and purification of PCR products (AMPure XP bead system). The libraries were analyzed for size distribution using an Agilent 2100 Bioanalyzer and were quantified using real-time PCR. For Oxford Nanopore Ultra-long sequencing, the libraries were prepared using the Ultra-Long DNA Sequencing Kit and using the standard protocol. The purified library was loaded onto primed Flow Cell and sequenced using a PromethION sequencer (Oxford Nanopore Technologies, Oxford, UK).

**RNA-seq and PacBio ISO-seq library preparation, construction and sequencing**

Total RNA was isolated from 6 distinct tissues (root, stem, tender leaves, flower, unripe fruits and ripe fruits) of XMM and DHP according to the instruction of Trizol method, respectively. After the integrity, quality and concentration of RNA, high quality RNA (2 μg, 300 ng/μL) was used to construct the Iso-Seq SMRTbell library via the SMRTbell® Express Template Prep Kit 3.0. RNA-seq sequencing libraries were generated using NEBNext® UltraTM RNA Library Prep Kit for Illumina® (NEB, USA) following manufacturer's instructions and index codes were added to attribute sequences to each sample. After cluster generation, the library preparations were sequenced on the Illumina Novaseq platform and 150 bp paired-end reads were generated.

**Whole-genome assembly**

First, raw sequencing reads were filtered adapter sequences, primers and other low-quality sequences using fastp v0.22.0 (Chen et al., 2018). Then, hifiasm v0.19.3 (Cheng et al., 2021) generated draft assembly. Subsequently, we aligned these contigs into *Fragaria vesca* 'Hawaii 4' (Zhou et al., 2023b) to assemble scaffolds and tune the chromosome ID and orientation using RagTag v2.1.0 (Alonge et al., 2022). Valid ligation products were detected from Hi-C data using HiC-Pro v3.1.0 (Servant et al., 2015), and only valid interaction pairs were retained for scaffolding of chromosomes using YaHS v1.2a.2 (Zhou et al., 2023a). EndHiC v1.0 (Wang et al., 2022) utilized the bin matrix results generated by HiC-pro as input data to create HiC heatmaps. In addition, several fragmented assemblies were generated to patch gaps and telomeres using Flye v2.9.2-b1786 (Kolmogorov et al., 2019) and Verkko v1.4.1 (Rautiainen et al., 2023) assemblers.

### Identification of telomere and centromere regions

Telomeres were detected utilizing the submit_telomere.sh available at https://github.com/VGP/vgp-assembly (Rhie et al., 2021). The monomers of centromeres (CEN17 and CEN18) were identified using TRASH (Wlodzimierz et al., 2023), and their distribution was confirmed through BLASTP v2.13.0+ (Ginestet, 2011). ModDotPlot v0.8.6 (Sweeten et al., 2024) was used to draw identity heatmaps. Ultimately, the centromere boundaries were determined by synthesizing information from the monomers density, transposable elements (TE) density, gene density, and Hi-C interaction map.

### Chromosome number analysis of XMM

The root tips of XMM which emerged after 3–4 days were collected and transferred into ice water for 24 h to accumulate metaphase cells. After which, they were fixed in a fresh standard solution of ethanol: acetic acid (3: 1, v/v) for 10 h at room temperature and stored at −20°C for at least 24 h. The seeding root-tip was digested in 1 mol/L hydrochloric acid for 8 min at 60°C. Thereafter, a chromosome slide was prepared using the conventional acetocarmine procedure. Chromosome images of the slides were captured using an OLYMPUS CX43 microscope. And the ploidy detection of XMM and DHP is estimated by flow cytometry analysis and *Fragra vesca* YW5AF7 (2n=2x=14) (Shulaev et al., 2011) was used as an internal control.

### Assembly quality assessments

Meryl v1.4 (Rhie et al., 2020) and GenomeScope v2.0 (Ranallo-Benavidez et al., 2020) were used to estimate chromosome ploidy, genome size and heterozygosity. Compleasm v0.2.5 (Huang and Li, 2023) was utilized to evaluate genome completeness using the eudicots_odb10 database. Genome accuracy was evaluated by aligning Long reads (HiFi and ONT) using minimap2 v2.24 (Li, 2018), and mapping rates and coverages were computed by SAMtools v1.17 (Danecek et al., 2021). LTR retrotransposons (LTR-RTs) are the predominant interspersed repeats that are poorly assembled in draft genomes, and they are used to evaluate assembly continuity through the LTR Assembly Index (LAI) v3.2 (Ou et al., 2018). We also assessed the accuracy, completeness, and phasing of assemblies generated by Merqury v1.3 (Rhie et al., 2020) using the Merqury 19-mer database, which integrates both Illumina PCR-free and HiFi reads.

**TE annotations**

Initially, homology-based and *ab initio* repeat library were constructed using famdb.py at https://github.com/Dfam-consortium/RepeatMasker and RepeatModeler v2.0.5 (Tarailo-Graovac and Chen, 2009), respectively. Subsequently, we combined two libraries and used The Extensive *de novo* TE Annotator (EDTA) v2.0.1 (Ou et al., 2019) for *de novo* TE predictions. The unknown Transposable Elements (TEs) were classified using DeepTE (Yan et al., 2020), and then queried within the assembled genome using RepeatMasker v4.1.5 (Flynn et al., 2020).

**Gene annotations and assessment**

A multi-strategy genomic annotation framework for XMM and DHP was implemented through systematic integration of transcriptomic evidence, homology inference, and *ab initio* predictions. RNA-Seq data were mapped to the XMM and DHP genomes using STAR v2.7.11a (Dobin et al., 2013) and assembled into transcripts using StringTie v.2.2.1 (Shumate et al., 2022). ISO-seq data were aligned, corrected, and high-confidence isoforms were defined using FLAIR v2.0.0 (Tang et al., 2020). TransDecoder v5.7.1 (https://github.com/TransDecoder/TransDecoder) identified candidate coding regions. In terms of protein homology, Rosaceae protein sequences were downloaded from the NCBI database and aligned against the XMM and DHP genome using miniprot v0.13-r262-dirty (Li and Valencia, 2023). BRAKER v3.0.7

(Stanke et al., 2008) was used to perform *ab initio* gene prediction using RNA-Seq, ISO-seq and homolog protein. The EVidenceModeler (EVM) v2.0.0 (Haas et al., 2008) software combined *ab initio* gene predictions and protein and transcript alignments into weighted consensus gene structures and the PASA pipeline v2.5.3 (Campbell et al., 2006) identified splicing variations and untranslated regions. We utilized BUSCO v5.4.5 (Manni et al., 2021) to estimate the completeness of genomic data in the annotated gene models, referencing the eudicots_odb10 database (Sep 2020, containing 2,326 sequences). eggNOG v2.1.12 (Jensen et al., 2007) is utilized for genome-wide functional annotation.

**Genome comparison between XMM and DHP**

The SyRI pipeline v1.6.3 (Goel et al., 2019) was used to detect structural variations between XMM and DHP genomes and plotsr v0.5.5 (Goel and Schneeberger, 2022) visualized structural similarities and rearrangements between the two genomes. The sequences which are not found in the reference genome were identified as presence/absence variants (PAVs) using ppsPCP v1.0 (Tahir Ul Qamar et al., 2019).

**Phylogenetic analyses**

Nonredundant protein sequences from 12 species (*Prunus armeniaca*, *Potentilla micrantha*, *Fragaria vesca*, *Rosa rugosa*, *Rosa wichuraiana*, *Rubus occidentalis*, *Rubus idaeus* 'Joan J', *Rubus argutus*, *Rubus chingii* Hu, *Rubus corchorifolius, Rubus hirsutus, Rubus eustephanos*) in the Genome Database for Rosaceae (Jung et al., 2019) were prepared for ortholog analyses. Orthogroups, orthologs, and the rooted species tree were inferred using OrthoFinder v2.5.5 (Emms and Kelly, 2019), and divergence times were calculated using the MCMCtree module from PAML v4.10.7 (Yang, 2007). The calibration times were obtained from the TimeTree database (www.timetree.org): for the genus species pairs of *R. idaeus* versus *R. occidentalis* (8.3 - 10.0 Mya), *Rubus* versus *Fragaria* (28.7 - 67.1 Mya), *Rosa wichuraiana* versus *Rosa rugosa* (3.3 -

41.0 Mya), *Fragaria vesca* versus *Potentilla micrantha* (20.4 - 51.2 Mya), *Rosa* versus *Fragaria* (21.1 - 57.8 Mya), and *Prunus* versus *Rosa* (49.2 - 77.1 Mya). Fianally, we used CAFE v5.0 (Mendes et al., 2021) to detect expansions and contractions of gene families.

**Transcriptome analysis**

Salmon v1.10.2 (Patro et al., 2017) was employed to generate highly accurate, transcript-level quantification estimates from RNA-seq data, while DESeq2 v1.44.0 (Love et al., 2014) was utilized to identify differentially expressed genes (DEGs) in ripe fruits of six high-sugar and six low-sugar groups. We conducted GO enrichment analysis utilizing clusterProfiler v4.12.6 (Xu et al., 2024).

**Measurement of total sugar content of *Rubus* fruits by UPLC-MS/MS**

Ripe fruits of ten *Rubus* species, named A1 to A8, XMM and DHP (Supplemental Figure 11A), were harvested from wood plant older than two years of age growing at Guizhou Botanical Garden and Northeast Agricultural University. The content of soluble sugar components was carried out by Guocangjian (http:// www.targetcrop.com/) based on the Sciex 4500 LC-MS/MS platform. Two biological replicates which consisted of five fruits from different plants were prepared for determination of total sugar content (fructose, glucose, sucrose and maltose).

Chemicals and reagents: ethanol, chloroform, acetonitrile (ACN) and formic acid (FA) were purchased from Merck (Darmstadt, Germany). Wastons pure water was used in all experiments. All of the standard and internal standard were purchased from Yuanye (Shanghai, China). The stock solutions of standards were prepared at the concentration of 0.3 mg/mL. All stock solutions were stored at 4 ℃.

Sample preparation and extraction: Fresh fruit materials were freeze dried in liquid nitrogen, and stored at -80℃ until needed. Then dried fruit materials were homogenized and powdered in a mill and then 30 mg of dried powder was extracted with 700 μL mixed solution of ethanol: $H_2O$ (80: 20). The extract was vortexed and under ultrasound for 30 minutes, then placed in 70℃ water bath for 2 hours. 700 μL chloroform was added and supernatants were collected after centrifugation. The supernatant was evaporated to dryness under nitrogen gas stream, then reconstituted in 100 μL mixed solution of ACN: $H_2O$ (75: 25). The solution was filtered through 0.22 μm filter for further LC-MS analysis.

HPLC conditions: The samples were analyzed using an HPLC-MS/MS system (HPLC ExionLC™ AC; MS Sciex Triple Quadrupole 4500). The HPLC analytical conditions were as follows: column was Inersil $NH_2$ column (5 μm, 4.6mm * 250 mm); solvent A was $H_2O$ (0.1 % FA); solvent B was ACN; flow rate was 1.0 mL/min with 75% B; column temperature was 40℃; injection volume was 10 μL.

MS conditions: Triple Quadrupole 4500 HPLC-MS/MS System, equipped with an ESI ion source, was operated in both positive and negative ion mode and controlled by Analyst 1.6.3 software (Sciex). The ESI source operation parameters were as follows: ion source, ESI-; ionspray voltage was 4500V, source temperature was 550℃; curtain gas (CUR) was set at 30 psi; the collision gas (CAD) was 9 psi. DP and CE for individual MRM transition was done with further DP and CE optimization. A specific set of MRM transitions were monitored for each period according to the sugars and sugar alcohols eluted within this period.

**Protein structure prediction**

The protein structure of *RhSTP7* and *RhSTP13* was predicted by AlphaFold 3 and visualized by PyMOL.

**Gene cloning, vector construction and RT-qPCR**

Tender leaves of XMM were sampled for RNA extraction according to the instructions of Polysaccharide Polyphenol Plant Total RNA Extraction Kit (TIANGEN, DP441). One microgram of total RNA was used for cDNA synthesis with a SPARKscript II RT Plus Kit (With gDNA Eraser) (Shandong Sparkjade Biotechnology Co., Ltd., AG0304-B). The cDNA product was diluted 10 times, and 1 µL was used as a template for amplifying the Coding DNA Sequence (CDS) of those two candidate genes and then integrated into over-expression construction JH23 via *Kpn*I and *Pac*I sites through homologous recombination (HR) using the ClonExpress Ultra One Step Cloning Kit (Vazyme), respectively, resulting in *RhSTP7*-OE and *RhSTP13*-OE constructs. As a negative control, the Cas9 fragment was also incorporated into JH23, resulting in Cas9-OE (JH23-Cas9). Primers used in this study are listed in Supplemental Table 12. All three constructs were transfected into *Agrobacterium* strain GV3101 and transiently infiltrated into *Rubus argutus* fruits at turning stage following the published protocols (Chatterjee et al., 2011). After three days, those injected-fruits were sampled for next study.

Total sugar content detection of the infiltrated-fruits was carried out by Sciex 4500 LC-MS/MS platform (Guocangjian). And total RNA of those injected-fruits expressing Cas9-OE, *RhSTP7*-OE and *RhSTP13*-OE constructs were extracted following above method. The cDNA product was synthesised by one microgram total RNA, and diluted 4 times, 1 µL was used as a template in RT-qPCR using QuantStudio 5 Systems (Applied Biosystems, Foster City, CA, USA) along with SYBR Green qPCR Mix (With ROX) (Shandong Sparkjade Biotechnology Co., Ltd., AH0104-B). The experiment was performed three times each with RNA extracted from newly samples (biological replicates). *RuEEF1A* was used as the housekeeping gene (Wu et al., 2021) for all RT-qPCR reactions and primers used for RT-qPCR are listed in Supplemental Table 12.

### The inversion verification of chromosome 6 centromere

To validate the inversion of chromosome 6 centromere of both XMM and DHP, specific primers (Supplemental Table 12) were designed for PCR reactions, which consisted of 15 μL PrimeSTAR GXL DNA Polymerase (Takara Bio, Japan), 1 μL of each upstream and downstream primer, 150 ng genomic DNA of XMM and $ddH_2O$ added to a total 30 μL volume. The PCR amplification was pre-denatured at 98℃ for 3 minutes and continued with 30 cycles of 30 seconds of 98℃ denaturation, 30 seconds of 58℃ annealing, and 1minute 30 seconds of 72℃ extensions. PCR products which were excised from the gel and purified via a NucleoSpin® Gel and PCR Clean-Up Kit (TaKaRa), then integrated into *pEASY*®-Blunt3 Cloning Vector (*pEASY*®-Blunt3 Cloning kit, TransGen Biotech) and performed by Sanger sequencing (Tsingke Biotech). The results of Sequence chromatograms were decoded using the SnapGene software (https://www.snapgene.com) (Supplemental Figure 7B). All Sanger-sequenced results were successfully aligned to the corresponding references with minimap2 (Li, 2018).

### Data visualization

Circos v0.69-8 (Krzywinski et al., 2009) was used to draw circular ideograms to depict genome structure variations. IGV v2.14.1 (Thorvaldsdóttir et al., 2013) was used to generate visualized tracks. Tree Visualization By One Table (tvBOT) (Xie et al., 2023) was used to display, manipulate, and annotate phylogenetic trees. Jvenn was used to generate Venn diagrams (Bardou et al., 2014). RectChr v1.36 was used to draw ideograms (https://github.com/BGI-shenzhen/RectChr). The R package ggplot2 v3.5.1 (Villanueva and Chen, 2019) was used to generate statistical representations and data graphics. The R package pheatmap v1.0.12 (Kolde and Kolde, 2015) generated RNA-seq heatmaps. HiCPlotter (Akdemir and Chin, 2015) was used to visualize Hi-C data.
